# An at-home Plant Physiology laboratory applied to dark-induced leaf senescence by college students and science teachers

**DOI:** 10.1101/2025.01.07.630824

**Authors:** Shejal Soumen, Alexander J. Kolstoe, Nora E. Duncan, Isaac P. Termolen, Amanda M. Solloway, Yan Lu

**Author notes:** **Correspondence** Yan Lu, Department of Biological Sciences, Western Michigan University, 1903 W Michigan Ave, Kalamazoo, MI 49008-5410, USA. Department of Plant & Soil Sciences, Texas Tech University, Lubbock, TX 79409, USA.

## Abstract

Dark-induced leaf senescence is an extreme example of leaf senescence induced by light deprivation. Prolonged dark treatments of individual leaves result in chlorophyll degradation, macromolecule catabolism, and reduction of photosynthesis. In this work, we described an at-home “Dark-induced Leaf Senescence” laboratory exercise for a junior-level undergraduate Plant Physiology course. To perform the dark-induced senescence assay on attached leaves, students may cover individual leaves of an outdoor plant with aluminum foils and record the leaf morphology with controlled vocabularies for ∼9 days. To perform senescence assays on detached leaves, the students may incubate detached leaves in various aqueous solutions (e.g., tap water, sucrose solution, alkali solution, and acid solution) either in the dark or under natural light, and then record the leaf morphology with controlled vocabularies for ∼9 days. This laboratory exercise provides hands-on opportunities for students to understand the relationships among sunlight, chlorophyll, and photosynthesis, in the comfort of students’ own homes. Specifically, it helps students to comprehend intrinsic and dark-induced leaf senescence mechanisms, the effects of sugars on leaf senescence, and the importance of optimal pH to plant health. This laboratory exercise can be adapted to support inquiry-based learning or be implemented in a middle or high school classroom.

## INTRODUCTION

Senescence is an energy-dependent, self-digesting process controlled by the interactions between environmental cues and developmental programs (Taiz et al., 2023). It is a universal characteristic in biological systems. According to the level of the senescing unit, plant senescence could be classified into: programmed cell death, organ senescence, and whole plant senescence (Taiz et al., 2023). All leaves, including those of evergreens (e.g., blue spruce), undergo senescence, in response to developmental factors (e.g., flowering and seeding), environmental factors (e.g., seasonal daylength and temperature changes), biotic stresses (e.g., pathogen attacks), or abiotic stresses (e.g., shading and wounding) (Taiz et al., 2023).

Intrinsic leaf senescence is a specialized form of programmed cell death, which permits remobilization of nutrients from source leaves to vegetative or reproductive sinks (Keskitalo et al., 2005). The earliest structural change during intrinsic leaf senescence is chloroplast breakdown (Taiz et al., 2023). Carbon fixation is thus replaced by the degradation and conversion of chlorophyll, proteins, and other macromolecules to exportable nutrients. Intrinsic leaf senescence is a normal developmental process (Kanojia et al., 2020).

Dark-induced leaf senescence is an extreme example of leaf senescence induced by shading (Sobieszczuk-Nowicka et al., 2018). Similar to intrinsic leaf senescence, dark-induced leaf senescence results in increased degradation of chlorophyll, disassembly of cellular elements (e.g., nucleic acids and proteins), and a loss of photosynthetic activity (Paluch-Lubawa et al., 2021). Dark-induced leaf senescence assays could be performed on whole plants, attached leaves, or detached leaves (Weaver and Amasino, 2001). This could be achieved by covering whole plants or individual leaves or by placing whole plants or detached leaves in the dark. Unlike whole plants or attached leaves, detached leaves are subjected to mechanical wounding and water-soaking (Iakimova and Woltering, 2018), as they need to be excised from the plant and kept in an aqueous solution. Mechanical wounds may act as additional entry points to detached leaves for substances present in the aqueous solution (Savatin et al., 2014).

During the social distancing imposed by COVID19 in Fall 2020 - Spring 2021, we developed an at-home laboratory topic – Dark-induced Leaf Senescence, for a junior-level undergraduate Plant Physiology course at Western Michigan University (WMU).

In this exercise, the students were asked to perform dark-induced leaf senescence assays on attached and detached leaves. For attached-leaf assays, the students may cover both sides of a few leaves (e.g., four) from a plant of their choice with aluminum foils. For detached leaf-assays, the students may excise some morphological and developmental similar leaves from a plant, keep them in aqueous solution, and place half of the leaves in the dark and the other half under natural light (e.g., by a window). The students were also asked to supplement the aqueous solution with sucrose, alkali (e.g., sodium bicarbonate/baking soda), or acid (e.g., acetic acid in vinegar and citric acid in lemon juice). Exogenous sugar treatments have been found to delay dark-induced leaf senescence in detached leaves (Wingler and Roitsch, 2008; Schippers et al., 2015; Li et al., 2020) and accelerate the senescence of detached leaves under light (Khudairi, 1970; Wingler et al., 2004; Wingler et al., 2006). A 6% sucrose solution was reported to be suitable for detached leaves or leaf segments (Li et al., 2020). Therefore, the students were asked to test whether supplying 6% sucrose to detached leaves delays or accelerate leaf senescence. Most plants thrive in the pH 6.0-7.0 (slightly acidic to neutral) range (Osman, 2018). The tap water in the Kalamazoo area has a pH of 7.0. Hence, the students were also asked to investigate the effect of pH on detached leaves by supplementing the aqueous solution with baking soda, which is sodium bicarbonate, or vinegar/lemon juice, which contains acetic acid or citric acid, respectively. A 6% baking soda solution has a pH of 8.0. A 6% vinegar solution has a pH of ∼3.2. A 6% lemon juice solution has a pH of ∼4.0. Before and during the treatments, the students were required to use controlled vocabulary to describe leaf morphology.

In Summer 2022, we modified this exercise slightly and showed it to 9 middle and high school science teachers from Southwest Michigan. They were participants of the Summer 2022 BIORETS (Research Experiences for Teachers Sites in Biological Sciences) program at WMU.

## LEARNING OBJECTIVES

The activities in this exercise should allow students to:

1. Understand leaf senescence mechanisms (e.g., intrinsic vs dark-induced leaf senescence) and the effects of sugars on leaf senescence.
2. Understand the importance of optimal pH to plant health.
3. Learn the basic techniques of dark-induced leaf senescence assays.
4. Use controlled vocabularies to record leaf morphology.

## MATERIALS AND METHODS

### Materials

In the lab manual (**Supplemental Material 1**), the students were provided with a list of materials used in this at-home laboratory exercise: outdoor plants with green leaves; aluminum foil; tap water; a measuring glass; eight glass/plastic jars/containers (e.g., Mason jars, jam jars, yeast jars, baby food jars, water glasses, small food storage containers made of clear plastics); a set of measuring spoons (e.g., one tablespoon); table sugar (i.e., sucrose); baking soda (i.e., sodium bicarbonate); vinegar, which contains acetic acid, or lemon juice, which contains citric acid; and a pair of scissors.

### Dark-Induced Leaf Senescence Assay with Attached Leaves

In the lab manual (**Supplemental Material 1**), the students were also provided with step-by-step instructions on how to perform dark-induced leaf senescence assays with attached and detached leaves. For the assay with attached leaves, the students chose four non-senescing green leaves from a plant and took pictures of each dark-treatment leaf, with at least one control leaf in the same picture. The eight leaves should be developmentally and morphologically similar. The students needed to use controlled vocabularies (**Table 1**) to record the initial leaf morphology (leaf color, presence or absence of necrotic spots or lesions) of the eight leaves (**Table 2**). The students then covered both sides of dark-treatment leaves with aluminum foils, secured the foils on the leaves by folding the foil near the tip and the base of the leaf inward, and labeled the leaves by tying a string on the petiole. If the students were concerned that the aluminum foil blocks the air and water vapor movements, they may replace the aluminum foil with black-colored fabric and secure the fabric with safety pins. The students also needed to label the four control leaves (e.g., by tying a string on each petiole). One day later, the students removed the aluminum foils and took pictures of each uncovered dark-treatment leaf, with at least one control leaf in the same picture. The students then recorded the leaf morphology of the eight leaves, re-covered the same four leaves with aluminum foils, and secured the foils. This process (morphology recording and imaging) may be repeated for 9 days for the dark-treated leaves to develop visible symptoms.

**Table 1.**
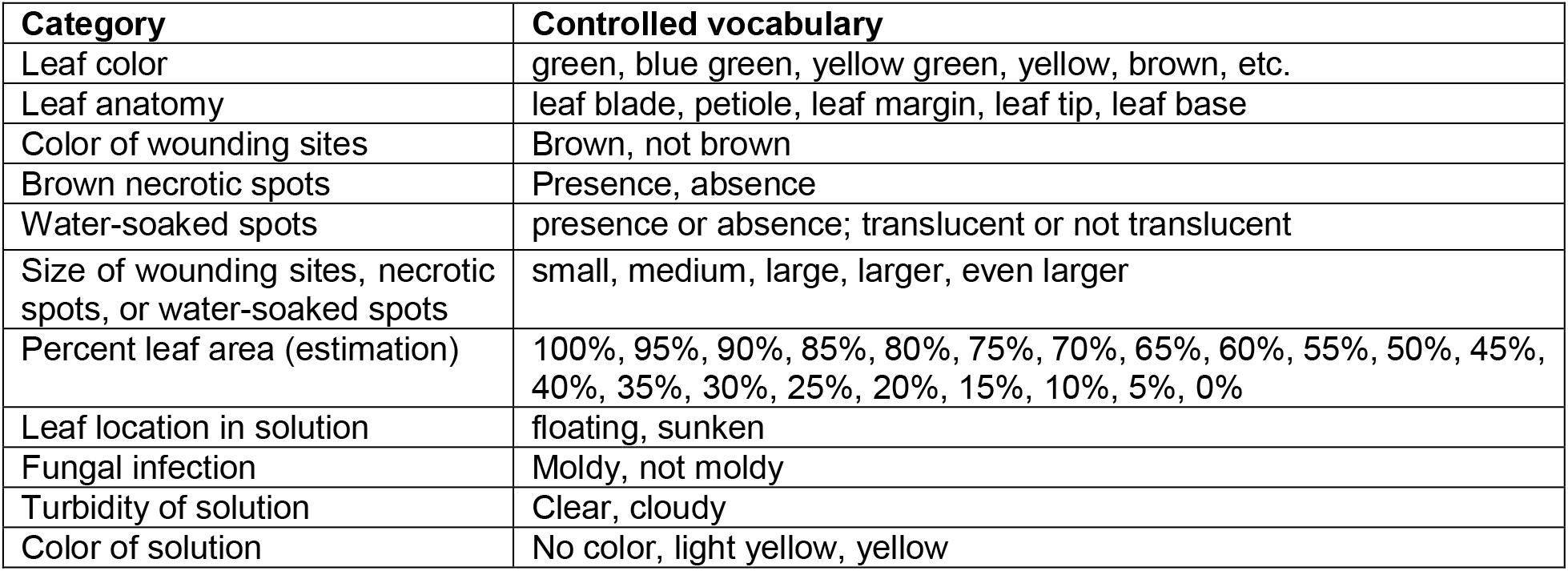
A list of controlled vocabularies to be used when recording leaf morphology.

| Category | Controlled vocabulary |
| --- | --- |
| Leaf color | green, blue green, yellow green, yellow, brown, etc. |
| Leaf anatomy | leaf blade, petiole, leaf margin, leaf tip, leaf base |
| Color of wounding sites | Brown, not brown |
| Brown necrotic spots | Presence, absence |
| Water-soaked spots | presence or absence; translucent or not translucent |
| Size of wounding sites, necrotic spots, or water-soaked spots | small, medium, large, larger, even larger |
| Percent leaf area (estimation) | 100%, 95%, 90%, 85%, 80%, 75%, 70%, 65%, 60%, 55%, 50%, 45%, 40%, 35%, 30%, 25%, 20%, 15%, 10%, 5%, 0% |
| Leaf location in solution | floating, sunken |
| Fungal infection | Moldy, not moldy |
| Turbidity of solution | Clear, cloudy |
| Color of solution | No color, light yellow, yellow |

**Table 2.**
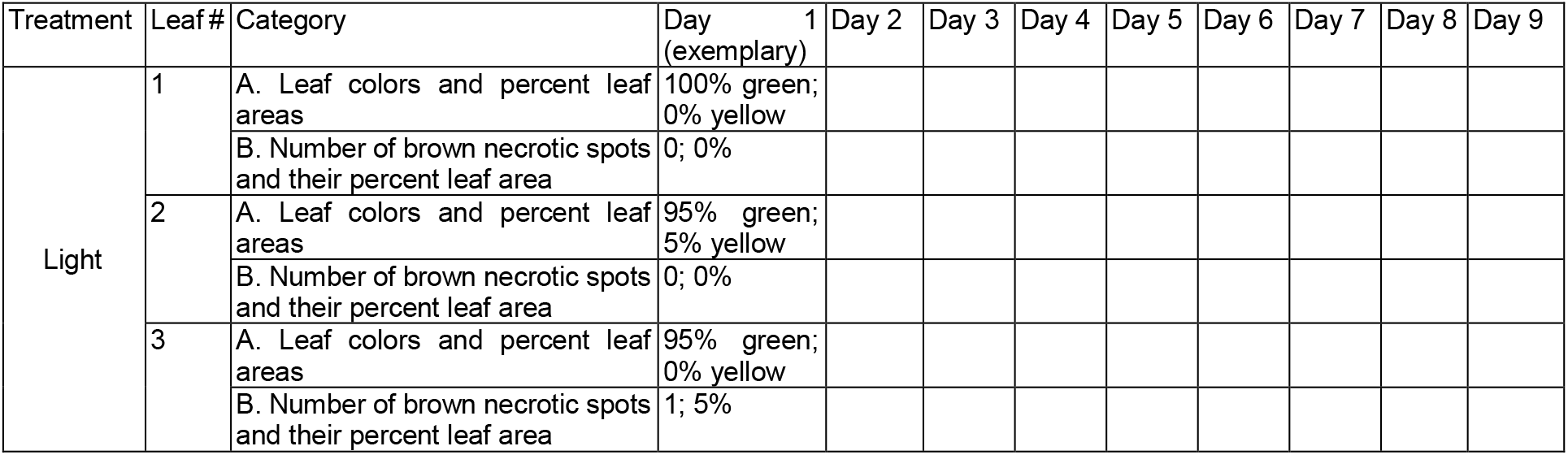
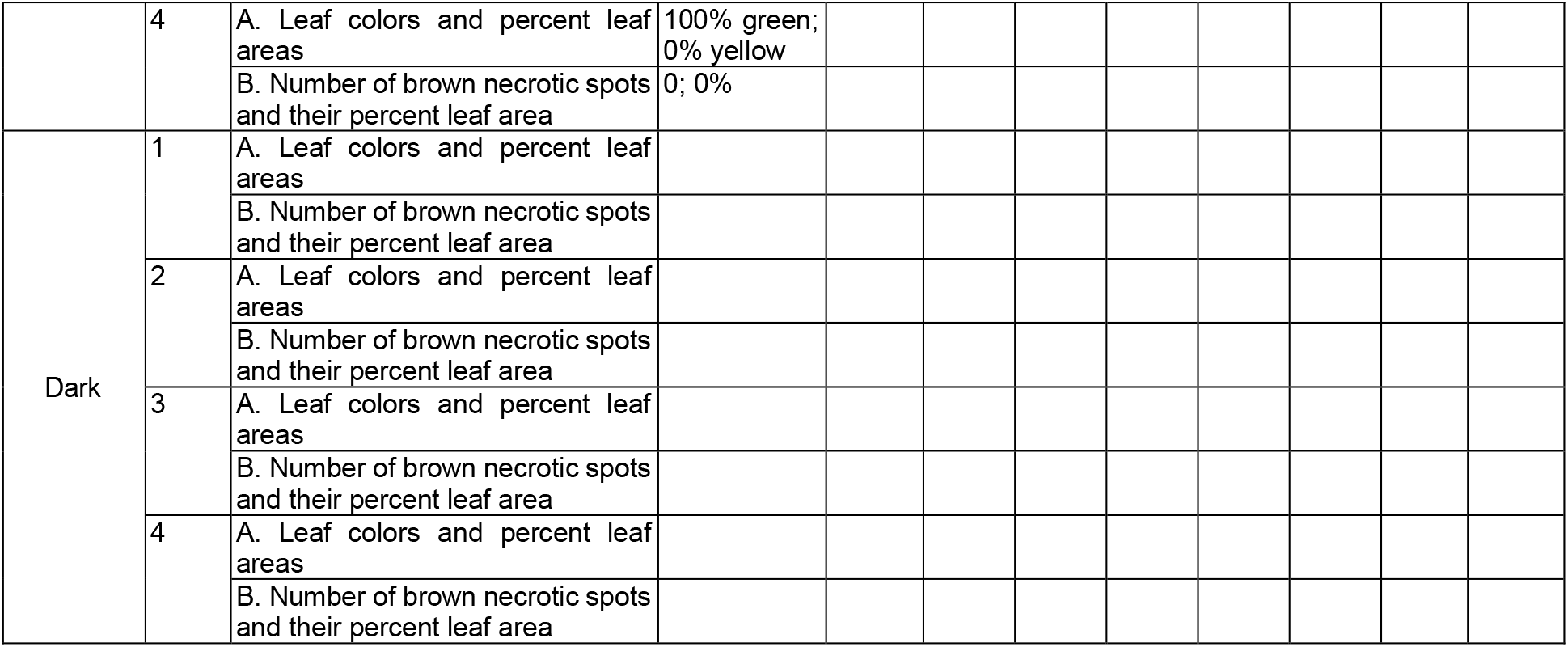
Daily morphology of dark-treated attached leaves.

### Dark-Induced Leaf Senescence Assay with Detached Leaves

For the assay with detached leaves, the students labeled 8 clear glass/plastic jars/glasses/containers with “H_2_O Light”, “H_2_O Dark”, “Sucrose Light”, “Sucrose Dark”, “Alkali Light”, “Alkali Dark”, “Acid Light”, and “Acid Dark”. For tap-water treatments, the students poured 1/2 cup (118 mL) of tap water into the jars labeled “H_2_O Light” and “H_2_O Dark”. For sucrose treatments, the students added 1 cup (237 mL) of tap water and 1 tablespoon (15 g) of table sugar (sucrose) into the jar labeled “Sucrose Light”, stirred with a stirring spoon to dissolve sucrose completely, and then transferred 1/2 cup of the resulting 6% sucrose solution into the jar labeled “Sucrose Dark”. After this, the students needed to wash the tablespoon, the stirring spoon, and the measuring glass with tap water and blot dry them with paper towels. For alkali treatments, the students added 1 cup (237 mL) of tap water and 1 tablespoon (15 g) of baking soda (sodium bicarbonate) into the jar labeled “Alkali Light”, stirred to dissolve the baking soda completely, and then transferred 1/2 cup of the resulting 6% baking soda solution into the jar labeled “Alkali Dark”. Again, the students needed to wash the tablespoon, the stirring spoon, and the measuring glass with tap water, and blot dry them with paper towels, after this step. For acid treatments, the students added 1 cup (237 mL) of tap water and 1 tablespoon (15 mL) of vinegar or lemon juice into the jar labeled “Acid Light”, stir to mix completely, and then transferred 1/2 cup of the resulting 6% acid solution into the jar labeled “Acid Dark”. After washing the tablespoon, the stirring spoon, and the measuring glass with tap water, the students set the eight jars aside (**Figure 1A**).

**Figure 1.**
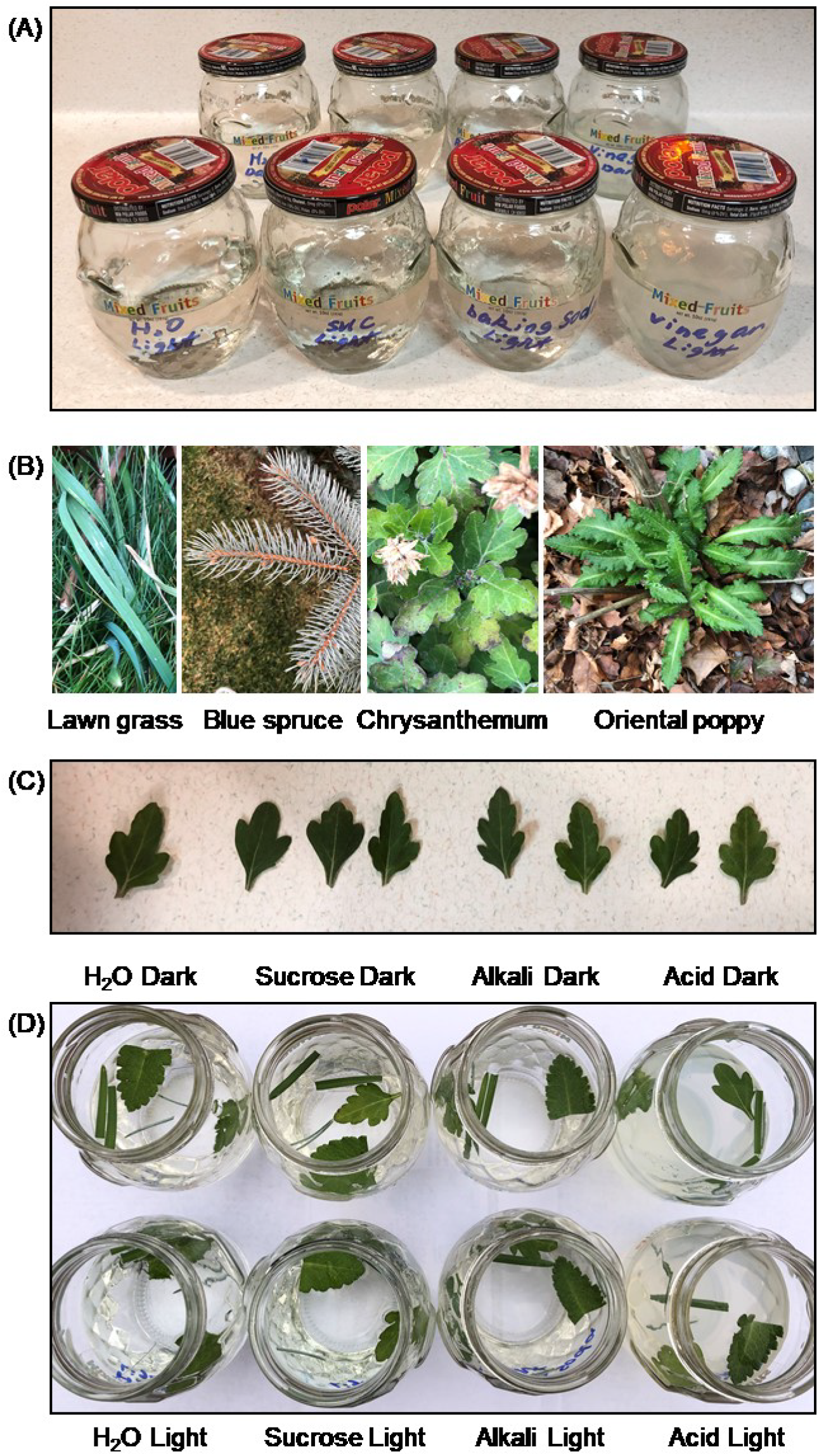
Dark-induced leaf senescence assay with detached leaves. (A) Eight jars with solutions. (B) Exemplary outdoor plants in winter 2020. (C) Arrange the leaves (e.g., Chrysanthemum leaves) according their size on a table. (D) Eight capless jars with detached leaves.

The students harvested ∼12 green leaves from the plant of their choice (exemplary plants in winter are shown in **Figure 1B**). These leaves should be non-senescing and developmentally and morphologically similar to each other. The students were also encouraged to include leaves from another plant in the assay, if they are interested. Multiple leaves could be incubated in each jar. The students then arranged the leaves according their size on a table, select 8 leaves that are non-senescing and most similar to each other developmentally and morphologically (**Figure 1C**), place one leaf per jar, and make sure all leaves face up. The students needed to use controlled vocabularies to record the morphology (leaf color, percentage of the leaf in that color, color of wounding sites, presence of brown necrotic spots and/or water-soaked spots, floating or sunken, etc.) of each leaf that goes into each jar, and the turbidity and color of each solution, in a table (see **Table 3**). After recording the morphology, the students took a group picture of the eight capless jars with leaves (**Figure 1D**) and placed the four jars labeled with “Light” under natural light (e.g., by a window) and the four jars labeled with “Dark” in the dark (e.g., in a drawer, cabinet, or closet). Capping the jars was optional during incubation. The students may repeat morphology recording and imaging every day for 9 days for the detached leaves to develop visible symptoms.

**Table 3.**
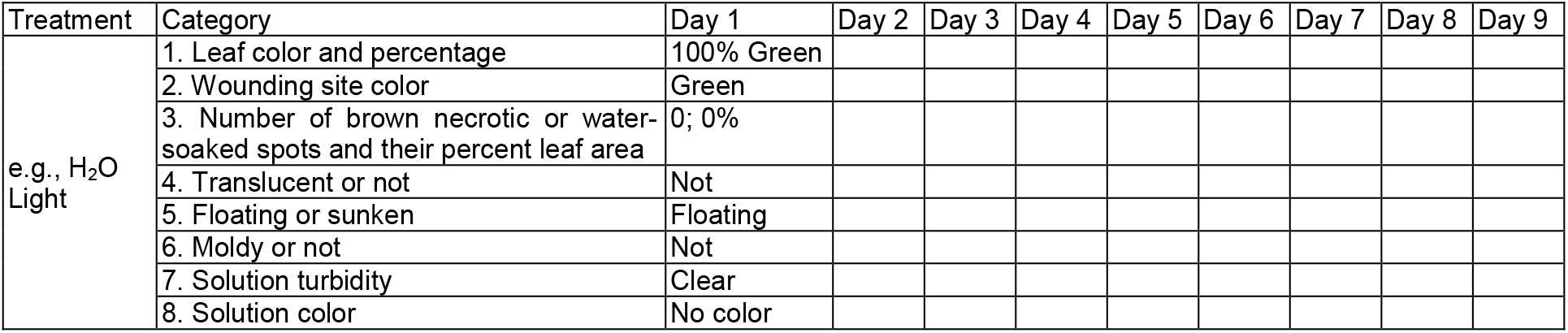
Daily morphology of dark-treated detached leaves.

## RESULTS

### Individually shaded, attached leaves displayed yellowing and senescence

We found that the attached leaves of various outdoor plants displayed yellowing and senescence after 7 days of shading with aluminum foils (**Figure 2**). Examples of such outplants include: common dandelions, day lilies, false bindweeds, hostas, prairie milkweeds, and yews. Performing the dark-induced leaf senescence assay may help students visually understand the mechanisms of intrinsic and dark-induced leaf senescence.

**Figure 2.**
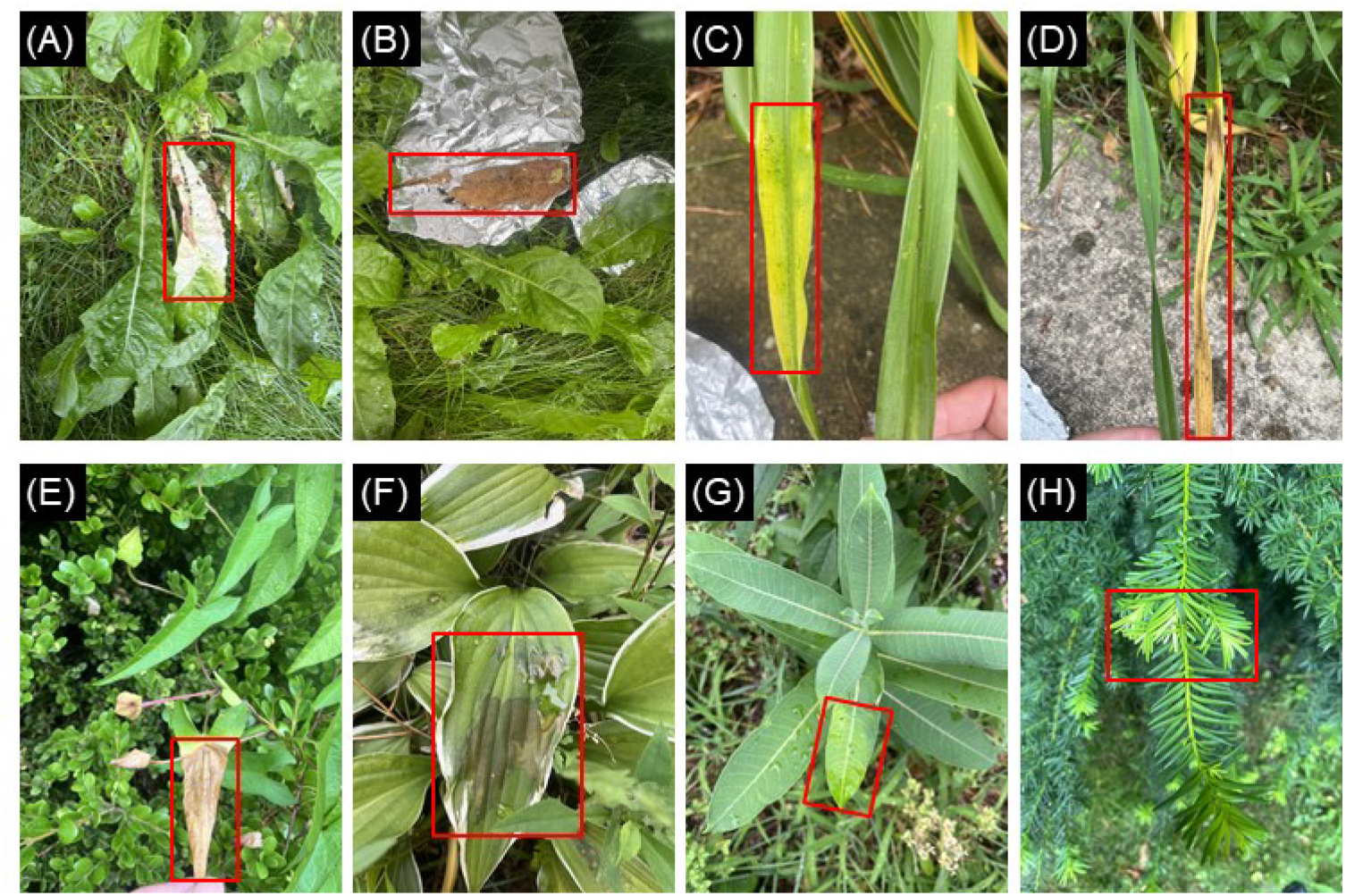
Examples of attached leaves or leaf sections of outdoor plants after 7 days of dark treatment in summer 2022. (A-B) Common dandelions. (C-D) Day lilies. (E) False bindweeds. (F) Hostas. (G) Prairie milkweeds. (H) Yews. Red rectangles indicate leaves or leaf sections covered with aluminum foils for 7 days.

### Detached leaves treated with “H_2_O + Dark” showed signs of senescence earlier than those treated with “H_2_O + Light”

We found that detached leaves placed in the dark in tap water showed signs of senescence earlier than those placed under nature light in the same tap water (**Figure 3**). After being incubated in tap water under nature light for 9 days (**Figure 3A**), the two lawn grass leaf sections and the poppy leaf section were still green. Although the Chrysanthemum leaf had three black necrotic spots, it still floated on top of water. The tap water was still clear. The leaves placed in the dark in the same tap water (**Figure 3B**) appeared less healthy. One of two lawn grass leaf sections turned yellow completely. In addition, the Chrysanthemum leaf and the poppy leaf section both sank to the bottom of the container, which is an extreme example of water-soaking. Furthermore, the tap water turned yellow, a sign of chloroplast destruction and chlorophyll leakage. These observations are consistent with the hypothesis that dark treatments result in leaf senescence.

**Figure 3.**
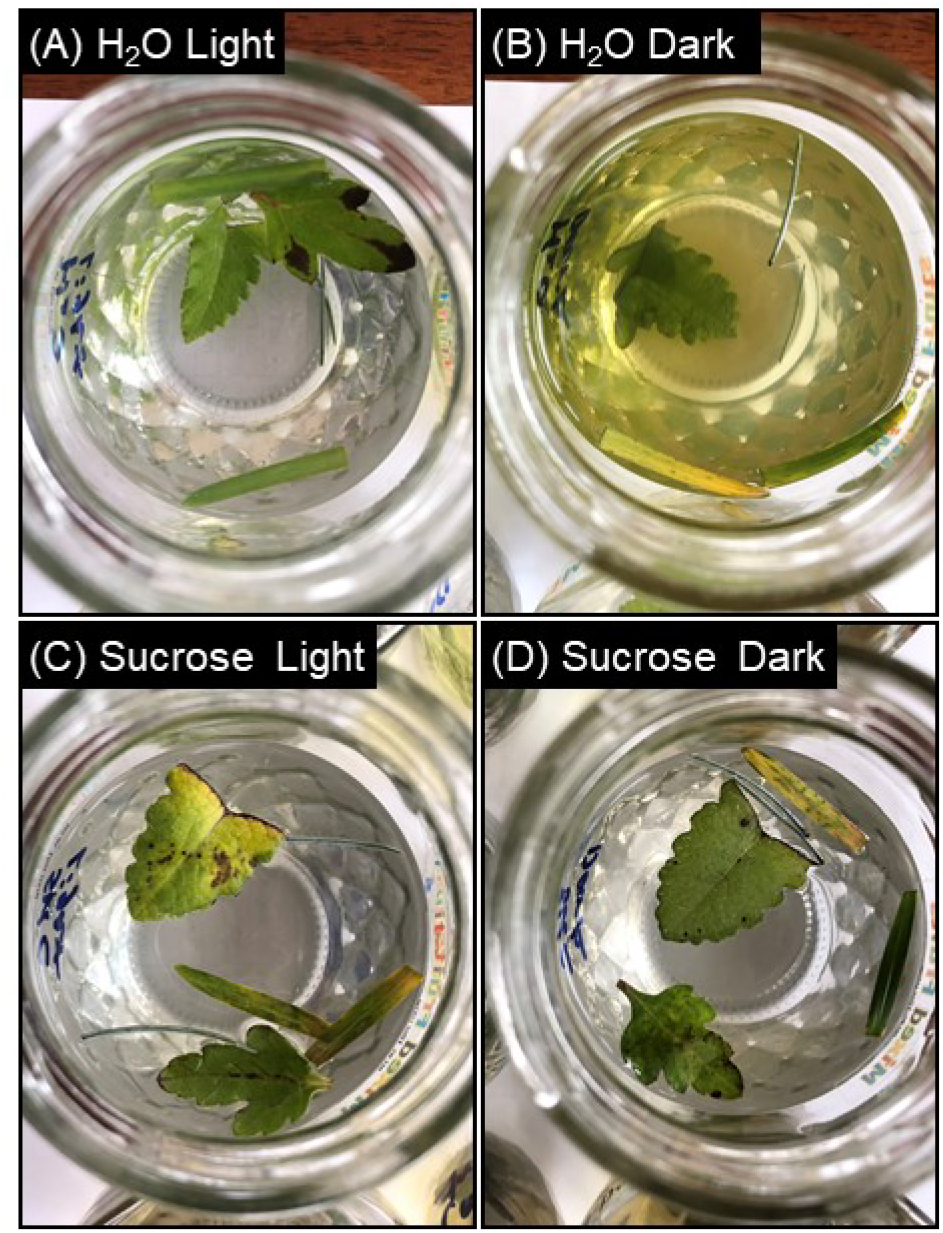
Detached leaves after 9 days of light or dark treatment in tap water or 6% sucrose.

### Detached leaves treated with “6% Sucrose + Light” showed signs of senescence earlier than those treated with “H_2_O + Light”

We found that detached leaves incubated in a 6% sucrose solution under nature light showed signs of senescence earlier than those incubated in tap water under the same nature light (**Figure 3**). After being incubated in 6% sucrose under natural light for 9 days (**Figure 3C**), the two lawn grass leaf sections, the Chrysanthemum leaf, and the poppy leaf section all turned yellow green and had many brown-to-black necrotic spots. These observations were initially surprising to the students because sucrose is a final product of photosynthesis and it can enhance plant growth. Interestingly, sugars also act as signaling molecules and regulate plant metabolism, development, and even senescence (Wingler et al., 2006). Sugar accumulations have been found to induce leaf senescence (Khudairi, 1970; Wingler et al., 2004).

### Detached leaves treated with “6% Sucrose + Dark” showed signs of senescence later than those treated with “H_2_O + Dark”

We found that detached leaves incubated in a 6% sucrose solution in the dark showed signs of senescence later than those incubated in tap water in the dark (**Figure 3**). After 9 days of dark treatment in 6% sucrose (**Figure 3D**), the Chrysanthemum leaf and the poppy leaf section were mostly green. As mentioned above, the Chrysanthemum leaf and the poppy leaf section subjected to 9 days of dark treatment in tap water sank to the bottom of the container and the tap water turned yellow (a sign of chloroplast destruction and chlorophyll leakage) (**Figure 3B**). These observations are consistent with the hypothesis that exogenous sugar may delay dark-induced leaf senescence in detached leaves (Wingler and Roitsch, 2008; Schippers et al., 2015; Li et al., 2020).

### Supplementing water with 6% baking soda caused damage to the detached leaves

We found that supplementing water with 6% baking soda caused damage to the detached leaves (**Figure 4**). After 3 days of incubation in a 6% baking soda solution under natural light, a large brown-to-black necrotic spot formed near the petiole of the Chrysanthemum leaf and the excision area of the poppy leaf section (**Figure 4C**). These two necrotic spots covered about 25% of the leaf area. After 5 days of incubation in a 6% baking soda solution under natural light, the necrotic spot covered about 50% of the poppy leaf section (**Figure 4D**). The two lawn grass leaf sections also developed necrotic spots near the excisions. Furthermore, the baking soda solution also became yellow, which is a sign of chloroplast destruction and chlorophyll leakage into the solution.

**Figure 4.**
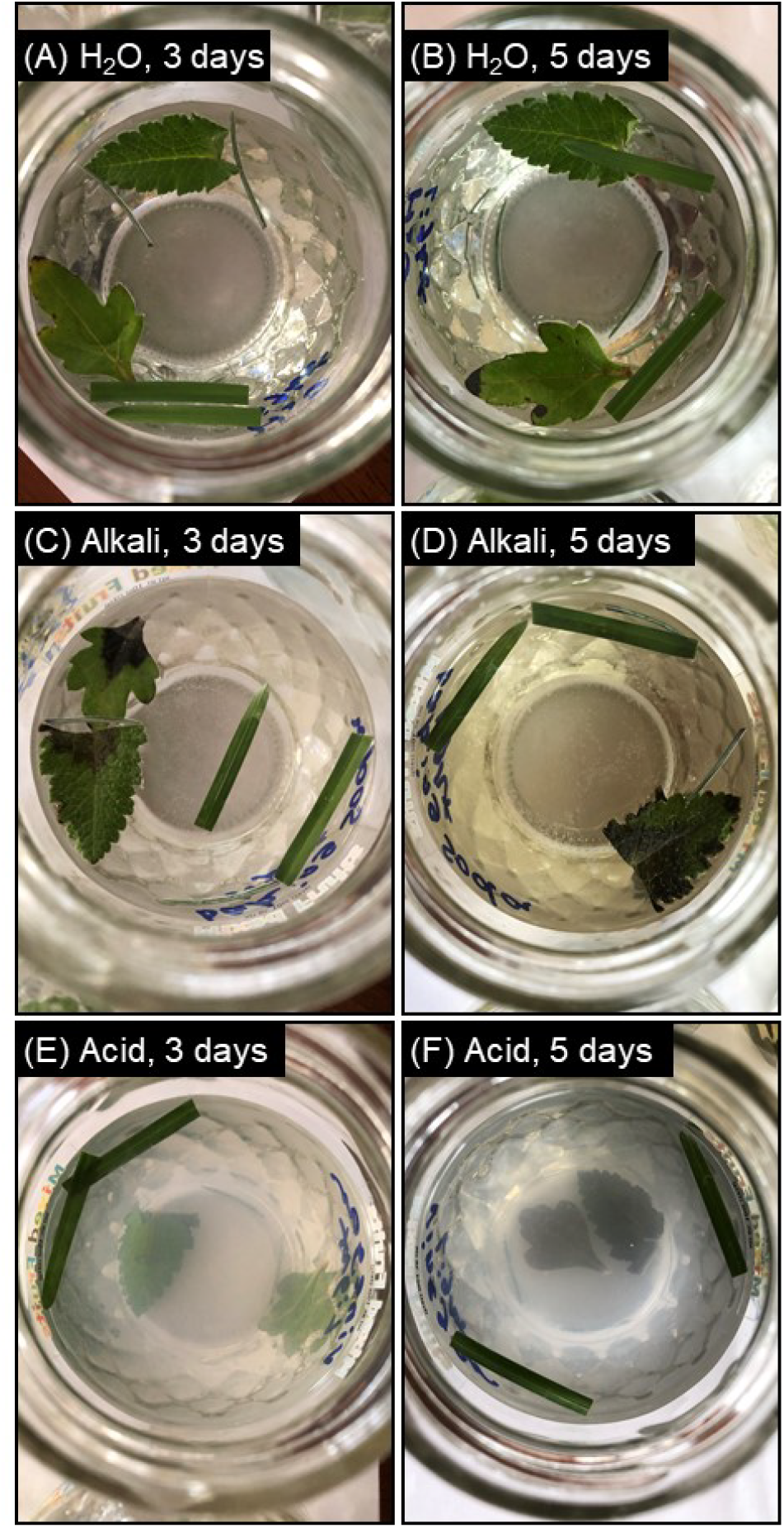
Detached leaves after being incubated in tap water (A-B), 6% baking soda (C-D), or 6% lime juice (E-F) under nature light for 3 (A, C, E) or 5 (B, D, F) days.

### Supplementing water with 6% vinegar or lemon juice caused damage to the detached leaves

We also found that supplementing water with 6% lemon juice caused damage to the detached leaves (**Figure 4**). After 3 days of incubation in a 6% lemon juice solution under natural light, the Chrysanthemum leaf and the poppy leaf section became translucent and sank to the bottom of the container (**Figure 4E**). After 5 days of incubation in a 6% lemon juice solution under natural light, the Chrysanthemum leaf and the poppy leaf section became dark brown (**Figure 4F**).

## DISCUSSION

### Effects of light on the senescence of attached and detached leaves

For many plant species, severe shading of leaves, especially when only applied to a part of the plant, results in rapid senescence (Liebsch and Keech, 2016). In this at-home laboratory exercise, after an individual leaf of an outdoor plant was covered with aluminum foils for 7 days, the leaf often turned yellow or even senesced (**Figure 2**). On the contrary, the control leaves not subjected to the dark treatment stayed green (**Figure 2**). Performing this at-home laboratory exercise allowed the students to visually understand that light deprivation is essential to the success of dark-induced senescence assays of both attached and detached leaves.

### Effects of sugars on the senescence of detached leaves

Exogenous sugar treatments have been found to accelerate the senescence of detached leaves under light but delay the senescence of detached leaves in the dark (Khudairi, 1970; Wingler et al., 2004; Wingler et al., 2006; Wingler and Roitsch, 2008; Schippers et al., 2015; Li et al., 2020). In this at-home laboratory exercise, detached leaves treated with 6% sucrose senesced earlier than those treated with tap water under nature light and senesced later than those treated with tap water in the dark (**Figure 3**). Therefore, performing this at-home laboratory exercise provided a hands-on opportunity for the students to understand the differential effects of sugars on the senescence of detached leaves under light or in the dark.

### Effects of pH on detached leaves

Most plants thrive in the pH 6.0-7.0 range (Osman, 2018). Treating plants with alkali or acidic solutions results in cell membrane leakage and water-soaking (Grant, 2024; Portland-Parks-and-Recreation, 2024). In this at-home laboratory exercise, detached leaves treated with 6% baking soda or 6% lemon juice showed signs of leaf damage (e.g., brown-to-black necrotic spots, translucent leaf coloration, sinking to the bottom) after 3 days of treatments and the symptoms worsened after 5 days of treatments (**Figure 4**). Therefore, carrying out this at-home laboratory exercise helped the students understand the importance of optimal pH to plant health.

### Connection between this at-home laboratory exercise and the corresponding lecture

Intrinsic leaf senescence is covered in one of the last four chapters of the “Plant Physiology and Development” textbook (Taiz et al., 2023) for the BIOS 3190 Plant Physiology course. During the online teaching of this chapter – “Plant Senescence and Developmental Cell Death”, students learned a number of related topics, such as the leaf senescence syndrome, the regulatory network of leaf senescence, and whole plant senescence. Therefore, having the students perform this at-home laboratory exercise near the end of the spring semester is complementary to and in sync with what the students have learned from the lectures. During the development stage of this laboratory module, we thought that performing dark-induced leaf senescence assays may help students understand the mechanisms of intrinsic and dark-induced leaf senescence. Indeed, one student stated in the final course evaluation that “I really enjoy doing the last lab at home as it was hands, which helped me learn and understand”.

### Completion rate of this at-home laboratory exercise

In Spring 2021, there were 11 undergraduate students enrolled in this junior-level BIOS 3190 Plant Physiology course. As a writing-intensive course, 32% of the overall grade came from lab reports and a total of nine lab reports were assigned. The first 8 lab topics were virtual and worth 16 points each (**Supplemental Material 2**). Dark-induced leaf senescence is the only at-home laboratory exercise and worth 40 points (**Supplemental Material 2**). Among the 11 students, 8 chose to complete this at-home exercise and submit a lab report about this laboratory topic. Therefore, the completion rate of this at-home exercise was 73%, similar to the average lab report completion rate of the 8 virtual topics (75%). This suggested that the extra work associated with the at-home laboratory exercise didn’t discourage the students from completing the lab and then submitting the lab report.

## POTENTIAL MODIFICATIONS

This laboratory exercise was developed during the COVID19 pandemic for undergraduate students to perform at home or in a classroom. If the students cannot find eight containers at home, they may drop the alkali or acid treatment. If the students have other class duties on certain days, they may opt out leaf morphology observation and photographing on these days. The students may also compare the images and morphology of detached leaves incubated in aqueous solutions with those attached to the plant to investigate the differences and similarities between detached and attached leaf senescence.

This at-home laboratory exercise can be easily adapted to an in-person classroom setting. For example, during the Summer 2022 BIORETS program, we had 9 middle and high school science teachers from Southwest Michigan performed this exercise in a classroom and it went very well. The images of attached leaf senescence assay shown in **Figure 2** were actually taken in Summer 2022.

This laboratory exercise, or part of this exercise, can also be simplified and implemented in a middle or high school science classroom as a hands-on activity for teaching photosynthesis. Chlorophyll is an essential component in photosynthesis. The simple and hands-on laboratory exercise described in this work may help students to visually understand the relationship among sunlight, chlorophyll, and photosynthesis. After performing this laboratory exercise during the Summer 2022 BIORETS program, some teacher participants remarked that they “could see how to implement it in their own classrooms”.

This laboratory exercise can also be adapted to support other pedagogical approaches, such as inquiry-based learning. For example, students may subject detached leaves to different concentrations of sucrose, baking soda, and vinegar/lemon juice and investigate whether different concentrations of sugars, alkali, or acids have differential effects on leaf health and senescence. Student may also place a set of detached leaves in the refrigerator and compare them with those incubated at room temperature. For an inquiry-based laboratory exercise, the students will be given a list of relevant references and will be asked to write a mini research proposal that contains an introduction and an experimental design section. In the introduction, the students are required to provide background information about their laboratory topic. In the experimental design section, the students are required to state their hypothesis, propose appropriate experiments, describe how to perform the experiments, list what equipment and materials they will need, define appropriate controls, explain what data they plan to collect, and clarify how they plan to analyze the data. After the students have finished the experiments and data collection, they will be asked to submit a lab report on this inquiry-based laboratory exercise, according to the grading criteria shown in **Supplemental Material 2**. Such inquiry-based laboratory exercises are expected to improve students’ motivation, critical thinking skills, and analysis skills (Buck et al., 2008; Díaz-Vázquez et al., 2012; Stefanou et al., 2013; Ambruso and Riley, 2022).

## CONCLUSION

In this work, we described an at-home laboratory exercise that was successfully implemented in an undergraduate Plant Physiology course. Materials needed for this exercise, such as aluminum foil, a measuring glass, glass/plastic containers, and a tablespoon, are readily available at students’ home. Therefore, performing this at-home laboratory exercise does not require shipping laboratory kits to students’ home. The activities involved this exercise should help students to: (1) understand leaf senescence mechanisms and the effects of sugars on leaf senescence; (2) understand the importance of optimal pH to plant health; (3) learn the basic techniques of dark-induced leaf senescence assays; (4) use controlled vocabularies to record leaf morphology. Activities included in this laboratory exercise are very flexible; students are encouraged to modify their experiments according to what they have at home. This laboratory exercise can also be adapted to support inquiry-based learning or be implemented as a hands-on activity for teaching photosynthesis in a middle or high school classroom.

## SUPPLEMENTAL MATERIALS

**Supplemental Material 1**. BIOS 3190 Plant Physiology Lab Manual on Dark-Induced Leaf Senescence.

**Supplemental Material 2**. BIOS 3190 Plant Physiology Point Distribution and Grading Criteria for Lab Reports.

## ACKNOWLEDGMENTS

The authors thank all the Spring 2021 BIOS 3190 Plant Physiology students and all the Summer 2022 BIORETS (Research Experiences for Teachers Sites in Biological Sciences) teacher participants at Western Michigan University (WMU). The authors also thank Mr. Christopher D. Jackson (WMU) for growth chamber management.

