## Supplemental Material 1 for "An at-home Plant Physiology laboratory applied to dark-induced leaf senescence by college students and science teachers"

**Supplemental Material 1.** BIOS 3190 Plant Physiology Lab Manual on Dark-Induced Leaf Senescence.

**Laboratory**

### *Dark-Induced Leaf Senescence*

**Learning Objectives: Understand leaf senescence mechanisms (e.g., intrinsic vs dark-induced leaf senescence) and the effects of sugars on leaf senescence; understand the importance of optimal pH to plant health; learn the basic techniques of dark-induced leaf senescence assays; and use controlled vocabularies to describe leaf morphology.**

**Note to students: You are given the opportunity to perform a dark-induced leaf senescence experiment in the comfort of your home. Although some images and data are provided in this manual, you are required to perform the experiments on your own and use the images and data you generated in your lab report.**

#### Introduction

Senescence is an energy-dependent, self-digesting process controlled by the interactions between environmental cues and developmental programs. It is a universal characteristic in biological systems. According to the level of the senescing unit, plant senescence could be classified into: programmed cell death, organ senescence, and whole plant senescence. All leaves, including those of evergreens (e.g., blue spruce), undergo senescence, in response to developmental factors (e.g., flowering and seeding), environmental factors (e.g., seasonal daylength and temperature changes), biotic stresses (e.g., pathogen attacks), or abiotic stresses (e.g., shading and wounding).

Intrinsic leaf senescence is a specialized form of programmed cell death, which permits remobilization of nutrients from source leaves to vegetative or reproductive sinks. The earliest structural change during intrinsic leaf senescence is chloroplast breakdown. Carbon fixation is thus replaced by the degradation and conversion of chlorophyll, proteins, and other macromolecules to exportable nutrients. Intrinsic leaf senescence is a normal developmental process.

Dark-induced leaf senescence is an extreme example of leaf senescence induced by shading. Dark-induced leaf senescence assays could be performed on whole plants, attached leaves, or detached leaves. This could be achieved by covering whole plants or individual leaves or by placing whole plants or detached leaves in the dark. Unlike whole plants or attached leaves, detached leaves are subjected to mechanical wounding and water-soaking, as they need to be excised from the plant and kept in an aqueous solution. Mechanical wounds may act as additional entry points to detached leaves for substances present in the aqueous solution.

#### Experiment Overview

In this exercise, you are going to perform dark-induced leaf senescence assays on attached and detached leaves. For attached-leaf assays, you may cover both sides of a few leaves (e.g., four) from a plant of your choice with aluminum foils. For detached leaf-assays, you may excise some morphological and developmental similar leaves from a plant of your choice, keep them in aqueous solution, and place half of the leaves in the dark and the other half under natural light (e.g., by a window). You will also supplement the aqueous solution with sucrose, alkali (e.g., sodium bicarbonate/baking soda), or acid (e.g., acetic acid in vinegar and citric acid in lemon juice).

Exogenous sugar treatment has been found to delay dark-induced leaf senescence in detached leaves and accelerate the senescence of detached leaves under light. A 6% sucrose solution was reported to be suitable for detached leaves or leaf segments. You are going to test whether supplying 6% sucrose to detached leaves delays or accelerates their senescence. Most plants thrive in the pH 6.0-7.0 (slightly acidic to neutral) range. The tap water in the Kalamazoo area has a pH of 7.0. You are going to investigate the effect of pH on detached leaves by supplementing the aqueous solution with baking soda, which is sodium bicarbonate, or vinegar/lemon juice, which contains acetic acid or citric acid, respectively. A 6% baking soda solution has a pH of 8.0. A 6% vinegar solution has a pH of ~3.2. A 6% lemon juice solution has a pH of ~4.0.

You are going to use controlled vocabulary to describe leaf morphology before and during the treatments. A list of controlled vocabulary is provided in the table below.

| **Category** | **Controlled vocabulary** |
|---|---|
| Leaf color | green, blue green, yellow green, yellow, brown, etc. |
| Leaf anatomy | leaf blade, petiole, leaf margin, leaf tip, leaf base |
| Color of wounding sites | brown or not brown |
| Brown necrotic spots | Presence or absence |
| Water-soaked spots | presence or absence; translucent or not translucent |
| Size of wounding sites, necrotic spots, or water-soaked spots | small, medium, large, larger, even larger |
| Percent leaf area (estimation) | 100%, 95%, 90%, 85%, 80%, 75%, 70%, 65%, 60%, 55%, 50%, 45%, 40%, 35%, 30%, 25%, 20%, 15%, 10%, 5%, 0% |
| Leaf location in solution | floating, sunken |
| Fungal infection | moldy or not moldy |
| Turbidity of solution | Clear or cloudy |
| Color of solution | No color, light yellow, or yellow |

**Definitions:**
**Leaf base:** the lowest part of a leaf blade that is near the petiole.
**Leaf blade:** the broad, expanded area of the leaf.
**Leaf margin:** the perimeter of the leaf between leaf tip and leaf base.
**Leaf tip:** the part of the leaf blade farthest from the petiole.
**Midvein:** the vein in the center of a leaf.
**Necrosis:** death that is directly caused by physical damage, toxins, or other external agents.
**Petiole:** the leaf stalk that join the leaf blade to the stem.
**Water-soaked spot:** a wet, dark, translucent appearance at the affected area. Incubation of detached leaves in an aqueous solution may result in water-soaked spots on the leaf blade. After prolonged incubation in an aqueous solution, the entire leaf may be water-soaked and sunken to the bottom. Prolonged incubation of photosynthetic tissues may also lead to chloroplast destruction and chlorophyll leakage into the solution. The resulting solution is light yellow or yellow colored.

#### Materials and Methods
##### Materials

Outdoor plants with green leaves (e.g., ground covers, herbaceous plants, shrubs, and trees)
Aluminum foil or light-proof (i.e., black-colored) fabric
Tap water
A measuring glass
Eight used and cleaned glass/plastic jars/containers (e.g., Mason jars, jam jars, yeast jars, baby food jars, water glasses, small food storage containers made of clear plastics; if you could not find eight containers at home, you may drop the alkali or acid treatment)
A set of measuring spoons (e.g., one tablespoon)

Table sugar (i.e., sucrose)
Baking soda (i.e., sodium bicarbonate)
Vinegar, which contains acetic acid, or lemon juice, which contains citric acid
A pair of scissors to harvest leaves

**Methods**

**Student Activity A: Dark-induced leaf senescence assay with attached leaves**

1. Choose four non-senescing green leaves from a plant of your choice, take pictures of each dark-treatment leaf, with at least one control leaf in the same picture. The eight leaves should be developmentally and morphologically similar. Record the leaf morphology (leaf color, presence or absence of necrotic spots or lesions) of the eight leaves with controlled vocabulary.
2. Cover both sides of a dark-treatment leaf with a piece of aluminum foil, secure the foil on the leaf by folding the foil near the tip and the base of the leaf inward, label the leaf by tying a string on the petiole. Repeat this process for three other dark-treatment leaves. If you are concerned that the aluminum foil blocks the air and water vapor movements, you may replace the aluminum foil with black-colored fabric and secure the fabric with safety pins.
3. Label the four control leaves by tying a string on each petiole.
4. One day (~24 hours) later, remove the aluminum foil, take pictures of each uncovered dark-treatment leaf, with at least one control leaf in the same picture. Record the leaf morphology of the eight leaves, re-cover the same four leaves with aluminum foils, and secure the foils. Repeat this process for 9 days.

| Treatment | Leaf # | Category | Day 1 (exemplary) | Day 2 | Day 3 | Day 4 | Day 5 | Day 6 | Day 7 | Day 8 | Day 9 |
|---|---|---|---|---|---|---|---|---|---|---|---|
| Light | 1 | A. Leaf colors and percent leaf areas | 100% green; 0% yellow | | | | | | | | |
| | | B. Number of brown necrotic spots and their percent leaf area | 0; 0% | | | | | | | | |
| | 2 | A. Leaf colors and percent leaf areas | 95% green; 5% yellow | | | | | | | | |
| | | B. Number of brown necrotic spots and their percent leaf area | 0; 0% | | | | | | | | |
| | 3 | A. Leaf colors and percent leaf areas | 95% green; 0% yellow | | | | | | | | |
| | | B. Number of brown necrotic spots and their percent leaf area | 1; 5% | | | | | | | | |
| | 4 | A. Leaf colors and percent leaf areas | 100% green; 0% yellow | | | | | | | | |
| | | B. Number of brown necrotic spots and their percent leaf area | 0; 0% | | | | | | | | |
| Dark | 1 | A. Leaf colors and percent leaf areas | | | | | | | | | |
| | | B. Number of brown necrotic spots and their percent leaf area | | | | | | | | | |
| | 2 | A. Leaf colors and percent leaf areas | | | | | | | | | |
| | | B. Number of brown necrotic spots and their percent leaf area | | | | | | | | | |
| | 3 | A. Leaf colors and percent leaf areas | | | | | | | | | |
| | | B. Number of brown necrotic spots and their percent leaf area | | | | | | | | | |

| | | |
|---|---|---|
| | 4 | A. Leaf colors and percent leaf areas |
| | | B. Number of brown necrotic spots and their percent leaf area |

**Student Activity C: Dark-induced leaf senescence assay with detached leaves**

1. Label 8 clear glass/plastic jars/glasses/containers with "$H_2O$ Light", "$H_2O$ Dark", "Sucrose Light", "Sucrose Dark", "Alkali Light", "Alkali Dark", "Acid Light", and "Acid Dark".
2. Measure 1/2 cup (118 mL) of tap water with a measuring glass, pour it into the jar labeled "$H_2O$ Light". Repeat this process for "$H_2O$ Dark".
3. Measure 1 cup (237 mL) of tap water, pour it into the jar labeled "Sucrose Light". Measure 1 tablespoon (15 g) of table sugar (sucrose), pour it into the same jar, and stir with a stirring spoon to dissolve sucrose. Pour 1/2 cup of the 6% sucrose solution into the measuring glass, transfer it into the jar labeled "Sucrose Dark". Wash the tablespoon, the stirring spoon, and the measuring glass with tap water. Dry the tablespoon with paper towels.
4. Measure 1 cup (237 mL) of tap water, pour it into the jar labeled "Alkali Light". Measure 1 tablespoon (15 g) of baking soda (sodium bicarbonate), pour it into the same jar, and stir to dissolve baking soda. Pour 1/2 cup of the 6% baking soda solution into the measuring glass, transfer it into the jar labeled "Alkali Dark". Wash the tablespoon, the stirring spoon, and the measuring glass with tap water. Dry the tablespoon with paper towels.
5. Measure 1 cup (237 mL) of tap water, pour it into the jar labeled "Acid Light". Measure 1 tablespoon of vinegar or lemon juice, pour it into the same jar, and stir to mix. Pour 1/2 cup of the 6% acid solution into the measuring glass, transfer it into the jar labeled "Acid Dark". Wash the tablespoon, the stirring spoon, and the measuring glass with tap water. Set the eight jars aside (see the figure below).

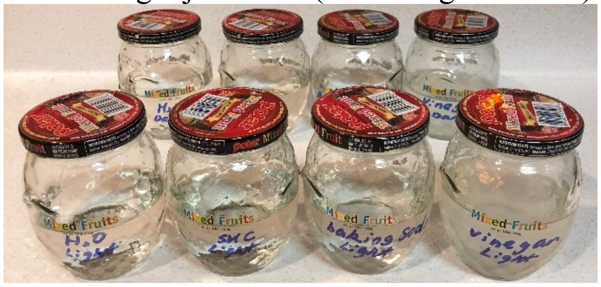

6. Harvest ~12 green leaves from the plant of your choice (exemplary plants in winter are shown in the figure below). These leaves should be non-senescing and developmentally and morphologically similar to each other. If you are interested in including leaves from another plant, please feel free to do so, multiple leaves could be incubated in each jar.

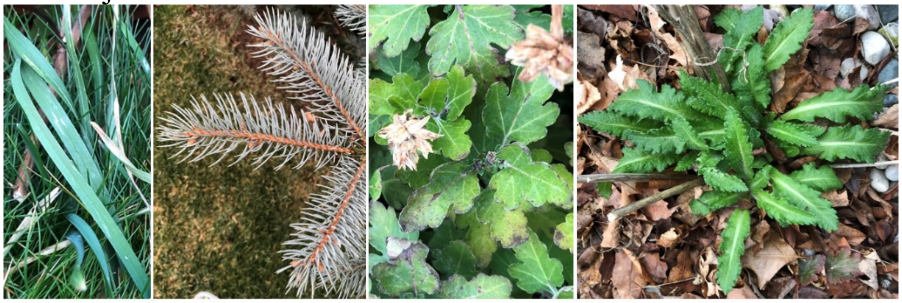

Lawn grass Blue spruce Chrysanthemum Oriental poppy

7. Arrange the leaves according their size on a table, select 8 leaves that are non-senescing and most similar to each other developmentally and morphologically (see the figure below). Place one leaf per jar and make sure all leaves face up.

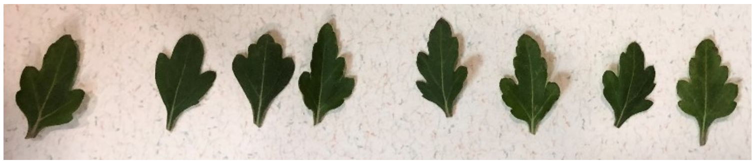

8. Record the morphology (leaf color, percentage of the leaf in that color, color of wounding sites, presence of brown necrotic spots and/or water-soaked spots, floating or sunken, etc.) of each leaf that goes into each jar, and the turbidity and color of each solution, in a table (see the table below), with controlled vocabulary.

| Treatment | Category | Day 1 | Day 2 | Day 3 | Day 4 | Day 5 | Day 6 | Day 7 | Day 8 | Day 9 |
|---|---|---|---|---|---|---|---|---|---|---|
| $H_2O$ Light | 1. Leaf color and percentage | 100% Green | | | | | | | | |
| | 2. Wounding site color | Green | | | | | | | | |
| | 3. Number of brown necrotic or water-soaked spots and their percent leaf area | 0; 0% | | | | | | | | |
| | 4. Translucent or not | Not | | | | | | | | |
| | 5. Floating or sunken | Floating | | | | | | | | |
| | 6. Moldy or not | Not | | | | | | | | |
| | 7. Solution turbidity | Clear | | | | | | | | |
| | 8. Solution color | No color | | | | | | | | |
| $H_2O$ Dark | 1. Leaf color and percentage | | | | | | | | | |
| | 2. Wounding site color | | | | | | | | | |
| | 3. Number of brown necrotic or water-soaked spots and their percent leaf area | | | | | | | | | |
| | 4. Translucent or not | | | | | | | | | |
| | 5. Floating or sunken | | | | | | | | | |
| | 6. Moldy or not | | | | | | | | | |
| | 7. Solution turbidity | | | | | | | | | |
| | 8. Solution color | | | | | | | | | |
| Sucrose Light | 1. Leaf color and percentage | | | | | | | | | |
| | 2. Wounding site color | | | | | | | | | |
| | 3. Number of brown necrotic or water-soaked spots and their percent leaf area | | | | | | | | | |
| | 4. Translucent or not | | | | | | | | | |
| | 5. Floating or sunken | | | | | | | | | |
| | 6. Moldy or not | | | | | | | | | |
| | 7. Solution turbidity | | | | | | | | | |
| | 8. Solution color | | | | | | | | | |
| Sucrose Dark | 1. Leaf color and percentage | | | | | | | | | |
| | 2. Wounding site color | | | | | | | | | |
| | 3. Number of brown necrotic or water-soaked spots and their percent leaf area | | | | | | | | | |
| | 4. Translucent or not | | | | | | | | | |
| | 5. Floating or sunken | | | | | | | | | |
| | 6. Moldy or not | | | | | | | | | |
| | 7. Solution turbidity | | | | | | | | | |
| | 8. Solution color | | | | | | | | | |
| Alkali Light | 1. Leaf color and percentage | | | | | | | | | |
| | 2. Wounding site color | | | | | | | | | |
| | 3. Number of brown necrotic or water-soaked spots and their percent leaf area | | | | | | | | | |
| | 4. Translucent or not | | | | | | | | | |
| | 5. Floating or sunken | | | | | | | | | |
| | 6. Moldy or not | | | | | | | | | |
| | 7. Solution turbidity | | | | | | | | | |

| | |
|---|---|
| | 8. Solution color |
| Alkali Dark | 1. Leaf color and percentage |
| | 2. Wounding site color |
| | 3. Number of brown necrotic or water-soaked spots and their percent leaf area |
| | 4. Translucent or not |
| | 5. Floating or sunken |
| | 6. Moldy or not |
| | 7. Solution turbidity |
| | 8. Solution color |
| Acid Light | 1. Leaf color and percentage |
| | 2. Wounding site color |
| | 3. Number of brown necrotic or water-soaked spots and their percent leaf area |
| | 4. Translucent or not |
| | 5. Floating or sunken |
| | 6. Moldy or not |
| | 7. Solution turbidity |
| | 8. Solution color |
| Acid Dark | 1. Leaf color and percentage |
| | 2. Wounding site color |
| | 3. Number of brown necrotic or water-soaked spots and their percent leaf area |
| | 4. Translucent or not |
| | 5. Floating or sunken |
| | 6. Moldy or not |
| | 7. Solution turbidity |
| | 8. Solution color |

9. Take a group picture and individual pictures of the eight capless jars with leaves.

**$H_2O$ Dark  Sucrose Dark  Alkali Dark  Acid Dark**

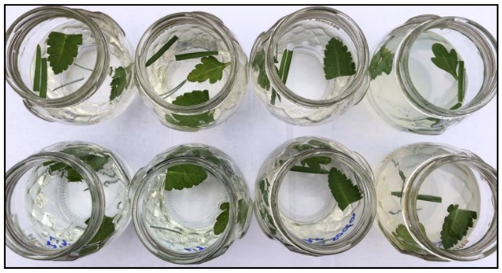

**$H_2O$ Light  Sucrose Light  Alkali Light  Acid Light**

10. Place the four jars labeled with “Light” under natural light (e.g., by a window) and the four jars labeled with “Dark” in the dark (e.g., in a drawer, cabinet, or closet). Capping the jars is optional during incubation.
11. Repeat steps 8-10 every day for 9 days.

**Results in the Lab Report:**
1. Use 1-2 sentences to summarize this laboratory.
2. Present data in tables and figures.
3. Report representative results with informative and descriptive sentences.

**Topics for Discussion in the Lab Report:**

1. Compare images and morphology of detached leaves in “$H_2O$ Light” and “$H_2O$ Dark”, leaves under which treatment show signs of senescence (yellowing, translucent leaf blades and brown necrotic or water-soaked spots) first? Use images you took yourself and leaf morphology you observed to support your answer (Exemplary images are shown below). Explain your observations.

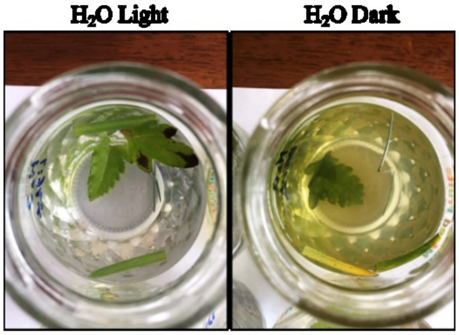

Day 9

2. Compare images and morphology of detached leaves in “$H_2O$ Light” and “Sucrose Light”, did supplementing water with 6% sucrose delay leaf senescence under light? Use images you took yourself and leaf morphology you observed to support your answer (Exemplary images are shown below). Explain your observations.

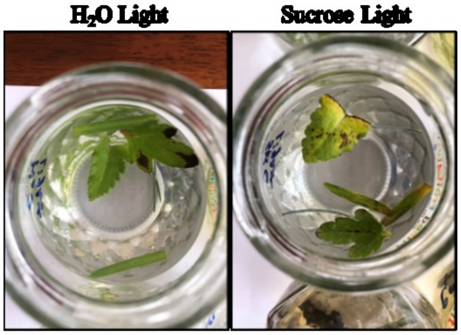

Day 9

3. Compare images and morphology of detached leaves in “$H_2O$ Dark” and “Sucrose Dark”, did supplementing water with 6% sucrose delay leaf senescence in the dark? Use images you took yourself and leaf morphology you observed to support your answer (Exemplary images are shown below). Explain your observations.

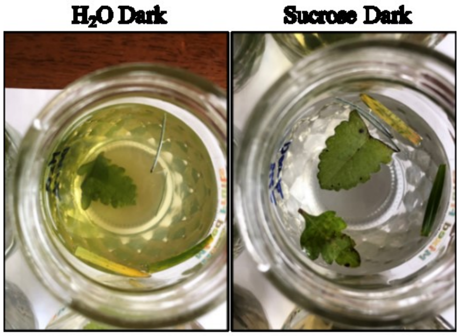

Day 9

4. Compare images and morphology of detached leaves in “$H_2O$ Light” and “Alkali Light”, did supplementing water with 6% baking soda cause leaf damage (translucent leaf blades and brown necrotic or water-soaked spots)? Use images you took yourself and leaf morphology you observed to support your answer (Exemplary images are shown below). Explain your observations. *(Hint: Consider the optimum pH range of plants.)*

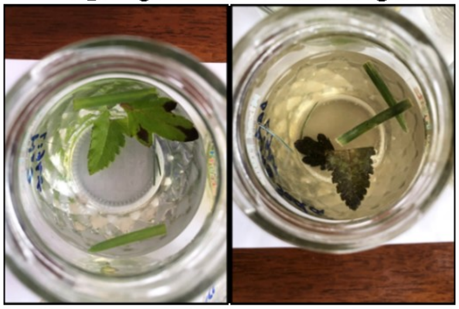

$H_2O$ Light | Alkali Light

Day 9

5. Compare images and morphology of detached leaves in “$H_2O$ Light” and “Acid Light”, did supplementing water with 6% vinegar or lemon juice cause leaf damage (translucent leaf blades and brown necrotic or water-soaked spots)? Use images you took yourself and leaf morphology you observed to support your answer (Exemplary images are shown below). Explain your observations. *(Hint: Consider the optimum pH range of plants.)*

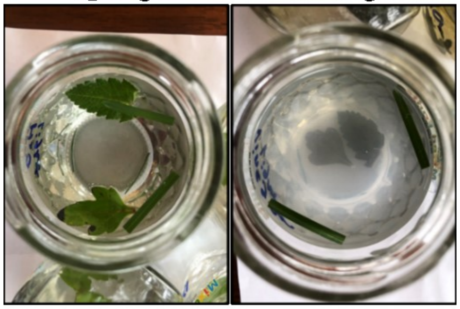

$H_2O$ Light | Acid Light

Day 5

6. Compare images and morphology of detached leaves in “Alkali Light” and “Acid Light”? Which treatment cause leaf damage (translucent leaf blades and brown necrotic or water-soaked spots) first? Use images you took yourself and leaf morphology you observed to support your answer (Exemplary images are shown below). Explain your observations. *(Hint: Consider pH difference between tap water and 6% baking soda solution and pH difference between tap water and 6% vinegar or lemon juice.)*

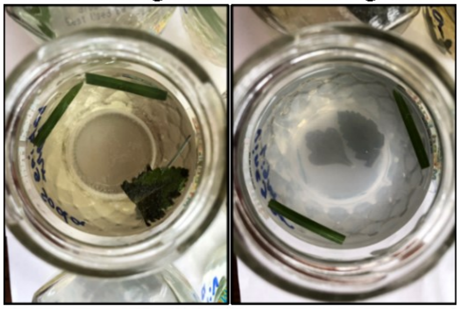

Alkali Light | Acid Light

Day 5

7. Compare images and morphology of detached leaves in “$H_2O$ Light” and attached leaves on the plant, which leaves show signs of senescence and/or damage first? Detached leaves or attached leaves? Use images you took yourself and leaf morphology you observed to support your answer (Exemplary images are shown below). Explain your observations, according to differences in plant species, experiment seasons, and indoor vs outdoor plants.

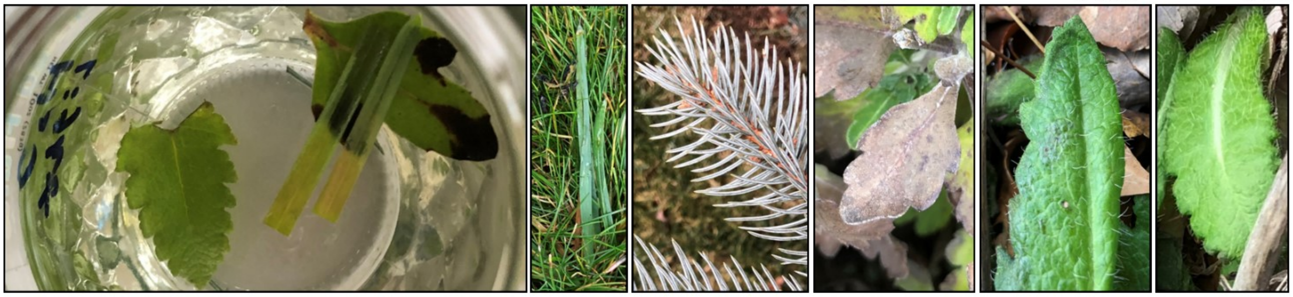

Day 12
