## Supplemental Material 2 for "An at-home Plant Physiology laboratory applied to dark-induced leaf senescence by college students and science teachers"

**Supplemental Material 2.** BIOS 3190 Plant Physiology Point Distribution and Grading Criteria for Lab Reports.

**Point distribution and grading criteria for lab reports are included here. Points will be deducted for not adhering to the expected formats and requirements as stated above and below.**

| | Point Distribution | | Grading Criteria |
|---|---|---|---|
| | Report 1-8 | Report 9 | |
| 1. Title | 0.5 | 0.5 | Use an informative, concise, and descriptive title |
| 2. Abstract | 2.0 | 5.5 | A brief but concise overview of the report, including the Introduction, Materials and Methods, Results, and the Discussion |
| 3. Introduction | 2.0 | 5.5 | Describe background information, as well as objectives, brief procedures, and importance of the experiment |
| 4. Materials and Methods | 2.0 | 5.5 | Address plant materials and growth conditions, experimental procedures, and data analysis approaches |
| 5. Results | 4.0 | 11.0 | Present data in tables and figures; report representative results with informative and descriptive sentences |
| 6. Discussion | 4.0 | 11.0 | Summarize results and discuss significance of the findings |
| 7. Literature cited | 0.5 | 1.0 | Include and correctly format sources of information |
| **TOTAL** | **15.0** | **40.0** | |
